# Subregion and sex differences in ethanol activation of cholinergic and glutamatergic cells in the mesopontine tegmentum

**DOI:** 10.1101/2023.11.08.566053

**Authors:** SM Mulloy, EM Aback, R Gao, S Engel, K Pawaskar, C Win, A Moua, L Hillukka, AM Lee

**Affiliations:** Graduate Program in Neuroscience, University of Minnesota, Minneapolis, MN, USA; Department of Pharmacology, University of Minnesota, Minneapolis, MN, USA

## Abstract

Ethanol engages cholinergic signaling and elicits endogenous acetylcholine release. Acetylcholine input to the midbrain originates from the mesopontine tegmentum (MPT), which is composed of the laterodorsal tegmentum (LDT) and the pedunculopontine tegmental nucleus (PPN). We investigated the effect of acute and chronic ethanol administration on cholinergic and glutamatergic neuron activation in the PPN and LDT in male and female mice. We show that ethanol selectively activates neurons of the PPN and not the LDT in male mice. Acute 4.0 g/kg and chronic 15 daily injections of 2.0 g/kg *i.p.* ethanol induced *Fos* expression in cholinergic and glutamatergic PPN neurons in male mice, whereas cholinergic and glutamatergic neurons of the LDT were unresponsive. In contrast, acute or chronic ethanol at either dose or duration had no effect on the activation of cholinergic or glutamatergic neurons in the MPT of female mice. Female mice had higher level of baseline activation in cholinergic neurons compared with males. We also found a population of co-labeled cholinergic and glutamatergic neurons in the PPN and LDT which were highly active in the saline- and ethanol-treated groups in both sexes. These findings illustrate the complex differential effects of ethanol across dose, time point, MPT subregion and sex.

## Introduction

Alcohol Use Disorder (AUD) is a major public health issue as 29.5 million people in the USA have an AUD and 140,000 annual deaths were attributed to alcohol-related causes in 2021 [1]. The prevalence of heavy alcohol use has increased over the past two decades and accelerated during the COVID pandemic [2,3]. The increased alcohol use that began during the pandemic is predicted to exacerbate alcohol-related morbidity and mortality rates [4,5]. Although there are pharmacological therapies for AUD (naltrexone, acamprosate, disulfiram), the neurobiology of alcohol needs to be better understood to identify new drug targets and assist the development of more effective treatments.

Alcohol (ethanol) and other misused drugs activate the ventral tegmental area (VTA) and cause dopamine release in the nucleus accumbens [6]. Voluntary ethanol consumption increases acetylcholine (ACh) levels in the VTA and dopamine levels in the nucleus accumbens in rats [7]. Inhibition of cholinergic signaling with intra-VTA administration of mecamylamine, a nicotinic acetylcholine receptor (nAChR) antagonist, blocks ethanol-induced increase in accumbal dopamine [8,9] and reduces ethanol intake [10]. Therefore, cholinergic signaling in the VTA is required for ethanol-induced dopamine release and ethanol consumption.

The only source of ACh input to the VTA originates from the mesopontine tegmentum (MPT), a brainstem region that is composed of two nuclei, the laterodorsal tegmentum (LDT) and the adjacent pedunculopontine tegmental nucleus (PPN) [11–13]. The MPT is primarily composed of cholinergic neurons but also contains glutamatergic and GABAergic neurons [14–16]. Cholinergic neurons form a continuum across the MPT and have numerous collaterals and projection targets including the VTA, substantia nigra, striatum, nucleus accumbens, habenula, interpeduncular nucleus, thalamus and medial prefrontal cortex [17]. MPT glutamatergic and GABAergic neurons also have extensive projection targets including the VTA and substantia nigra [14,17,18].

Prior studies have shown that nicotine injections increase c-Fos expression in the PPN in male rats [19], and modulation of the PPN and LDT via lesions or pharmacological inhibitors alter consumption and seeking for nicotine and cocaine [20–22]. Although ACh in the VTA is required for ethanol-induced increase in dopamine levels [8,9], and this ACh originates from the cholinergic MPT, relatively little is known about how ethanol affects neurons in the MPT. Prior studies have not examined the effect of ethanol on MPT subregions, cholinergic or glutamatergic cell types or across sex. In this study, we investigated both acute and chronic ethanol effects on the activation of cholinergic and glutamatergic neurons in the PPN and LDT in male and female mice. We found that cholinergic and glutamatergic PPN neurons from male mice expressed *Fos* after acute high dose ethanol and chronic low dose ethanol, whereas cholinergic and glutamatergic neurons of the LDT were unresponsive. In contrast, acute or chronic ethanol at any dose or time point had no effect on the activation of cholinergic or glutamatergic neurons in the MPT of female mice, and instead produced a trend for a reduction in cholinergic neuronal activity after ethanol administration. These data highlight the sex differences in ethanol-induced activation of cholinergic and glutamatergic neurons in the MPT.

## Methods

### Animals and drug administration

Male and female C57BL/6J adult 56-day old mice (The Jackson Laboratory) were grouped housed and acclimated to our facility for 5-7 days after arrival. Mice were habituated to 3 days of handling and daily intraperitoneal (*i.p.*) saline injections before experimental treatment. Ethanol was diluted to 20% (v/v) in injectable saline. For acute administration, mice received a single *i.p.* injection of 2.0 or 4.0 g/kg ethanol, or equivalent volume of saline. For chronic administration, mice received daily *i.p.* injections of 2.0 g/kg ethanol or saline. Mice were euthanized and brains collected 2 hours after the last injection.

### Immunohistochemistry (IHC)

Brains were sectioned at 30μm and coronal LDT and PPN sections were collected. Sections were blocked with 5% normal donkey and alpaca serum solution, and incubated overnight at 4°C with anti-ChAT (choline acetyltransferase) goat polyclonal antibody (1:500, Invitrogen #PIPA518518) and anti-c-Fos rabbit polyclonal antibody (1:1000, Abcam #ab190289). The secondary antibodies were anti-goat donkey Alexafluor488 (1:400, JIR #705-546-147) and anti-rabbit alpaca Alexafluor594 (1:400, JIR #611-585-215). Sections were mounted using Prolong Gold+DAPI and stored in the dark at 4°C.

### RNAscope *in situ* hybridization

Brains were sectioned at 16μm and processed using RNAscope fluorescent *in situ* hybridization Multiplex-V1 (Advanced Cell Diagnostics, Inc., Bio-Techne) according to manufacturer recommendations. Target probes purchased from Advanced Cell Diagnostics were Mm-*Chat*-c1 (choline acetyltransferase, #408731), Mm-*Fos*-c2 (#316921), and Mm-*Slc17a6*-c3 (Vglut2, vesicular glutamate transport 2, #319171). Fluorescence amplification 4A was used, and resulting probes were GFP-ChAT, mcherry-Fos, and cy5-Vglut2. All slides were stained for 4’,6-Diamidino-2-phenylindole (DAPI), mounted with Prolong Gold Antifade mounting media and stored at 4°C in the dark.

### Immunohistochemistry and RNAscope image analysis

All images were captured using a Keyence BZ-X700 microscope. Each region of interest (ROI) consisted of a left or right section from the LDT or PPN. IHC images were quantified by at least 2 experimenters blinded to treatment conditions. Blinded images were hand-counted and the total number of cholinergic neurons in each ROI as well as the number of cholinergic neurons co-labelled with c-Fos were quantified by each experimenter. The percent of cholinergic neurons expressing c-Fos (ChAT+cFos+ / all ChAT+) was calculated for each ROI. Blinded images with ROIs that had less than 20 or more than 250 cholinergic neurons (ChAT+) were excluded (15 of 263 ROIs collected). Of these remaining ROIs, we excluded 7 based on statistical outliers (outside of upper or lower limits of interquartile range as defined as 1.5x interquartile range above or below the quartile and when the Z-score was outside of ± 2.68).

For RNAscope, ROIs were analyzed in an automated, unbiased manner based on our prior work [23,24]. Each image was loaded into MATLAB Computer Vision Toolbox (MathWorks; Natick, MA) for image registration followed by ROI selection, which were based on *Chat* expression in the LDT or PPN. Registered ROI image sets were exported from MATLAB to CellProfiler (www.cellprofiler.org, RRID:SCR_007358) [25–28]. The background was subtracted and intensity information was split into HSV color space and converted to grayscale. Cell areas were automatically identified in an unbiased manner based on DAPI expression and the Ostu thresholding method. CellProfiler was then used to relate the signal punctum to cell area for each probe in an unbiased manner to generate the number of probe puncta per cell in each ROI. Data were imported into MATLAB, where the the probability that a cell was positive for a target probe was calculated using a binomial distribution test comparing the background probe expression and the puncta expression within the cell area, with an α of 0.01 and a Bonferroni correction using the number of cells in that ROI. The percent of cells per ROI that were positive for a single, two or three target probes of interest was then calculated and averaged across ROIs per treatment group. The level of expression of *Fos* transcript in a cell was calculated as the total pixel volume of *Fos* transcript divided by the total pixel volume of a target cell X 100, which was calculated for all target cells in each ROI, and reported as average % *Fos* transcript. ROIs that had fewer than 20 or more than 300 cholinergic neurons were excluded (34 out of 228 ROIs collected). Of these remaining ROIs, we excluded 17 based on statistical outliers (outside of upper or lower limits of interquartile range as defined as 1.5x interquartile range above or below the quartile and when the Z-score was outside of ± 2.68).

### Statistical analysis

To evaluate sex differences, a fixed effects mixed model with a random effect for the mouse variable was used to determine differences between sex, region, and treatment (JMP pro 17, SAS Institute Inc.). If multiple comparisons testing was warranted, a corrected Student’s *t*-test pairwise comparison was performed. As we have previously reported sex differences in cholinergic signaling in mice [29], we expected and observed significant sex differences in neuronal activation by ethanol, thus all subsequent data was analyzed separately by sex. For IHC data, the percent of activated cholinergic neurons (ChAT+, c-Fos+) was analyzed using a nested one-way ANOVA within each sub-region (PPN or LDT). For RNAscope data, the average percent of activated neurons per ROI and the average percent amount of *Fos* transcript in each target cell per ROI was compared using a nested t-test or nested one-way ANOVA (Prism 10, GraphPad, La Jolla, CA). Nested analyses and random effects were used to account for non-independence of the data points. All data are reported as mean±SEM.

## Results

### Immunohistochemical identification of cholinergic neurons in the MPT

The MPT subregions, PPN and LDT, are defined by cholinergic neuron expression around the superior cerebellar peduncle (**Fig. 1**). We first determined if the number of cholinergic neurons (ChAT+ cells) in the PPN or LDT differed by sex or was affected by ethanol treatment. There was no difference in the number of cholinergic neurons in the PPN between groups injected with saline, 2.0 g/kg ethanol, a rewarding dose in mice [30], or 4.0 g/kg ethanol, a sedating and aversive dose in mice [31], across sex as there was no sex by ethanol interaction and no main effects of sex or ethanol treatment (F_sexXtreatment_(2, 18.9)=1.275, *P*=0.30; F_sex_(1, 19.9)=2.993, *P*=0.10; F_treatment_(2, 18.9)=0.500, *P*=0.61, **Fig. S1**). There was no difference in the total number of cholinergic neurons in the PPN between male and female mice chronically treated with saline or 2 g/kg ethanol as there were no main effects or interactions between sex, treatment or day (all *P*>0.05). In the LDT, the number of cholinergic neurons was also not different between male or female mice acutely treated with saline, 2.0 or 4.0 g/kg ethanol (F_sexXtreatment_(2, 16.5)=0.605, *P*=0.56; F_sex_(1, 17.0)=0.0001, *P*=0.99; F_treatment_(2, 16.5)=0.464, *P*=0.64), or chronically treated with saline or 2.0 g/kg ethanol (all *P*>0.05, **Fig. S1**). Overall, these data show that the number of cholinergic neurons was consistent between sex, MPT subregion and across ethanol treatment dose and duration. Therefore, the observed differences in the percent of activated cholinergic neurons after ethanol are not confounded by differences in the total number of cholinergic neurons.

**Figure 1.**
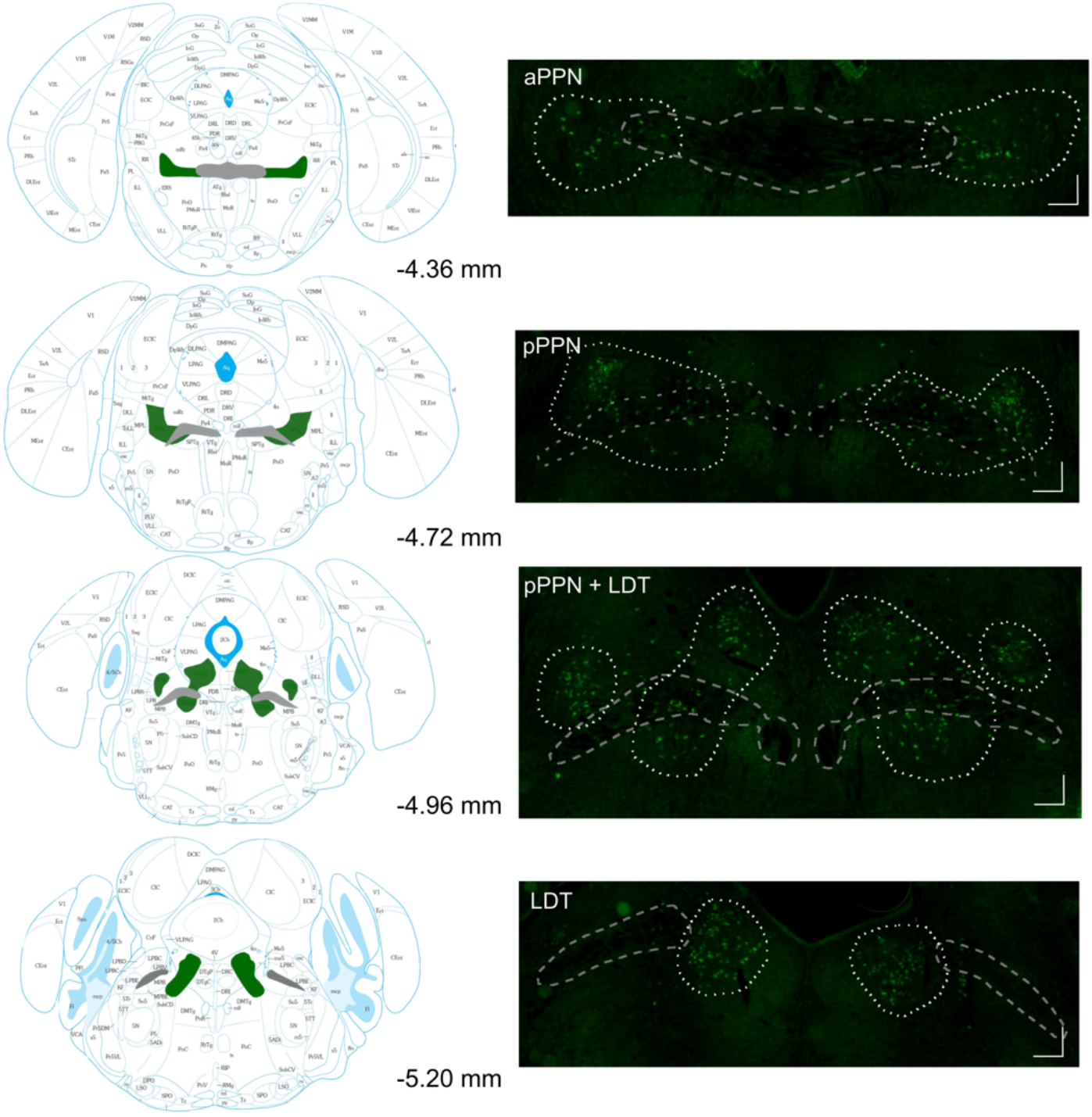
Outline of the MPT subregions PPN and LDT. The subregions of the cholinergic MPT, the anterior pedunculopontine tegmentum (aPPN), posterior pedunculopontine tegmentum (pPPN) and laterodorsal tegmentum (LDT) are defined by the distribution of cholinergic neurons (green) around the superior cerebellar peduncle (grey). Atlas and bregma coordinates adapted from Paxinos & Franklin, 2008. Scale bar = 100 μm.

### Acute ethanol effects in MPT cholinergic neurons

We determined the effect of acute administration of saline, 2.0 or 4.0 g/kg ethanol on cholinergic PPN and LDT neuron activation by probing for c-Fos expression using IHC. We found a sex difference in the percent of activated cholinergic neurons (ChAT+ and c-Fos+ cells) in the PPN after acute ethanol injection. There was a significant interaction between sex and ethanol dose (F_sexXdose_(2, 19.5)=7.324, \*\**P*=0.004; F_sex_(1, 21.0)=5.95, *P*=0.02; F_dose_(2,19.5)=0.188, *P*=0.83). Multiple comparisons testing with corrected Student’s *t*-tests showed a difference in the percent of activated cholinergic neurons between male and female mice administered saline (*P*=0.02) and after administration of 4.0 g/kg ethanol (*P*=0.01). These data show that cholinergic neurons are differentially active in male and female mice at baseline and differentially activated after an injection of a high, sedating dose of ethanol. Based on the baseline sex difference in cholinergic neuron activation, males and females were analyzed separately for all analyses.

In the PPN of male mice, the percent of cholinergic neuron activation was significantly different across groups (nested one-way ANOVA, F(2,10)=17.10, *P*=0.0006) and a Tukey’s multiple comparisons test showed that one 4.0 g/kg ethanol injection increased the percent of activated cholinergic neurons in the PPN by 2.3-fold over saline (\*\*\**P*<0.001 for 4.0 g/kg vs. saline) and 1.7-fold over the 2.0 g/kg ethanol dose (\*\**P*<0.01 4.0 vs 2.0 g/kg ethanol; **Fig. 2a**). There was no significant difference in the percent of activated PPN cholinergic neurons between saline and 2.0 g/kg ethanol (*P*=0.31) with saline resulting in 16.8 ± 2.0%, 2.0 g/kg ethanol in 23.2 ± 2.8%, and 4.0 g/kg ethanol in 38.9 ± 2.7% activated cholinergic neurons. In the LDT of male mice, saline resulted in 10.4 ± 3.0%, 2.0 g/kg ethanol in 18.6 ± 5.2%, and 4.0 g/kg ethanol in 24.8 ± 3.1% activated cholinergic neurons (**Fig. 2b**). There was no significant difference in the percent of activated cholinergic neurons between the treatment groups in the LDT (F(2,8)=1.130, *P*=0.37), highlighting a subregion difference in cholinergic neuron activation.

**Figure 2.**
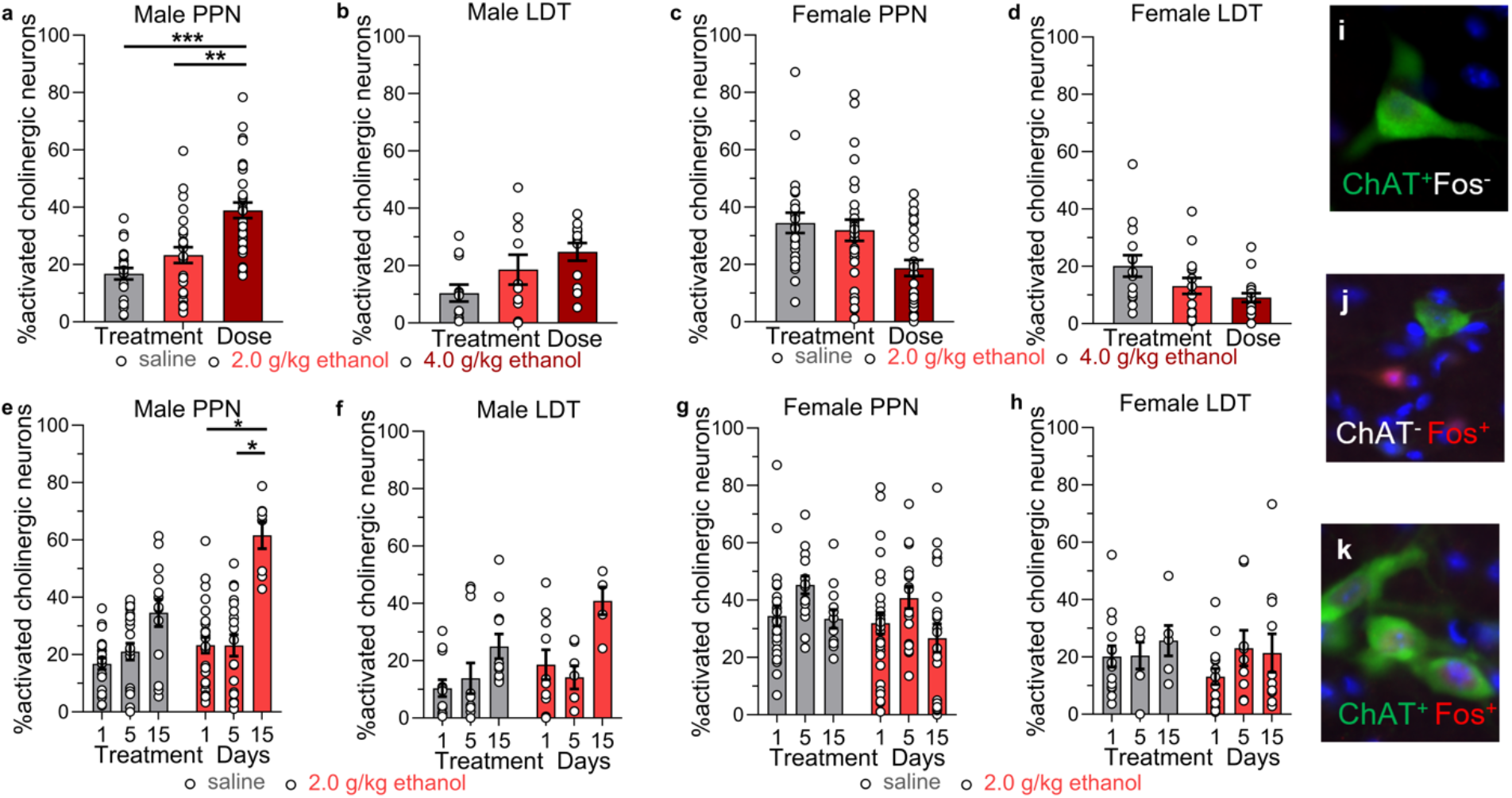
Acute and chronic ethanol activation of cholinergic neurons in the PPN and LDT. **a)** Ethanol (4.0 g/kg *i.p.*) increased the percent of c-Fos+ cholinergic neurons in the PPN but not the **b)** LDT of male mice measured with IHC. Tukey’s multiple comparisons test:\*\*\**P*<0.001, 4.0 g/kg vs saline; \*\**P*<0.01 4.0 g/kg vs 2.0 g/kg ethanol. PPN: *n*=21-33 ROI from 4-5 mice/group; LDT: *n*=10-12 ROI from 3-4 mice/group. **c-d)** Acute ethanol did not increase c-Fos+ cholinergic neurons in female PPN or LDT. PPN: *n*=24-29 ROI from 4 mice/group; LDT: *n*=14-17 ROI from 4-5 mice/group. **e)** Chronic ethanol (2.0 g/kg/day for 15 days) increased the percent of c-Fos+ cholinergic neurons in the PPN in male mice. Tukey’s multiple comparison test: \**P*=0.04, days 1 vs 15, \**P*=0.049, days 5 vs 15. PPN: *n*=8-27 ROI from 2-4 mice/ethanol group, *n*=14-22 ROI from 3-4 mice/saline group; LDT: *n*=5-10 ROI from 2-3 mice/ethanol group, *n*=11-12 ROI from 3-4 mice/saline group. **f-h)** Chronic ethanol did not increase c-Fos+ cholinergic neurons in male LDT or in female PPN or LDT. PPN: *n*=18-29 ROI from 3-4 mice/ethanol group, *n*=12-24 ROI from 2-4 mice/saline group; LDT: *n*=9-15 ROI from 3-4 mice/ethanol group, *n*=6-14 ROI from 2-4 mice/saline group. **i-k)** Example images of cells positive and negative for ChAT and c-Fos. Data expressed as mean ± SEM.

In the PPN of female mice, acute saline treatment resulted in 34.5 ± 3.5%, 2.0 g/kg ethanol in 31.9 ± 3.7%, and 4.0 g/kg ethanol in 18.7 ± 2.8% activated cholinergic neurons (**Fig. 2c**). Ethanol dose did not significantly change the percent of activated cholinergic neurons, although there was a trend for decreased cholinergic neuron activation after 4.0 g/kg ethanol compared with saline in the PPN (**Fig. 2c**; F(2,9)=1.563, *P*=0.26). In the LDT of female mice, there was also no significant effect of treatment group, with saline resulting in 20.1 ± 3.8%, 2.0 g/kg ethanol resulting in 13.2 ± 2.8%, and 4.0 g/kg ethanol in 9.1 ± 1.5% cholinergic neuron activation (**Fig. 2d**; F(2,9)=1.155, *P*=0.36). Together, these data indicate that sex and MPT subregion are important factors in the responsiveness to ethanol, with male PPN cholinergic neurons activated by 4.0 g/kg ethanol, whereas there was no significant change in cholinergic neuron activation in the LDT of male mice or in the PPN or LDT of female mice.

### Effect of chronic rewarding ethanol treatment on MPT cholinergic neurons

We next determined the effect of 5 or 15 daily injections of 2.0 g/kg ethanol or saline and compared them with the single 2.0 g/kg or saline administration group to obtain a time course of 1, 5, and 15 days of treatment. In the PPN of male mice, one day of saline resulted in 16.8 ± 2.0% activated cholinergic neurons, 5 days of saline in 21.1 ± 2.9% and 15 days of saline in 34.6 ± 4.8% activated cholinergic neurons. There was no significant difference in the percent of activated cholinergic neurons after saline treatment across these time points (F(2,8)=1.488, *P*=0.28) indicating that chronic saline treatment did not change the percent of activated cholinergic neurons in the PPN (**Fig. 2e**). A single 2.0 g/kg ethanol injection resulted in 23.3 ± 2.8%, 5 days of injections in 23.2 ± 3.8% and 15 days of injections in 61.7 ± 4.7% activated cholinergic neurons in the PPN. There was a significant difference between groups (F(2,6)=5.792, *P*=0.04) and a Tukey’s multiple comparison test showed a significant difference in cholinergic neuron activation between days 1 and 15 (\**P*=0.04) and between days 5 and 15 (\**P*=0.049, **Fig. 2e**). There was no difference between days 1 and 5 (*P*=0.99). These data show that 2.0 g/kg ethanol activates PPN cholinergic neurons after 15 days of daily injections but not after 1 or 5 days of injections.

In the LDT in male mice, there was no significant difference in the percent of activated cholinergic neurons in mice injected with 1, 5, or 15 days of saline (F(2,8)=0.7500, *P*=0.50) or between mice injected with 1, 5, or 15 days of 2 g/kg ethanol (F(2, 5)=2.678, *P*=0.16), with 18.6 ± 5.2% activated cholinergic neurons after 1 day, 14.2 ± 4.0% after 5 days and 40.9 ± 4.8% after 15 days of ethanol injections (**Fig. 2f**).

In contrast to male mice, female mice showed no effect of chronic 2.0 g/kg ethanol or saline injections on the percent of activated cholinergic neurons in the PPN (ethanol F(2,7)=0.5736, *P*=0.59; saline F(2,5)=0.5065, *P*=0.78, **Fig. 2g**) or in the LDT (ethanol F(2,7)=0.2965, *P*=0.75; saline F(2,5)=0.2507, *P*=0.79, **Fig. 2h**). These data show that cholinergic neurons in the PPN, but not in the LDT, of male mice are activated after 15 days of daily injections of a rewarding 2.0 g/kg ethanol dose but not after 1 or 5 days of ethanol. In contrast, female mice show no activation of PPN or LDT cholinergic neurons after the same chronic ethanol treatment.

### Identification of neuronal cell types in the MPT with RNAscope

The MPT is heterogenous and contains glutamatergic neurons in addition to cholinergic neurons. We compared the activation of glutamatergic versus cholinergic neurons after 15 days of daily 2 g/kg ethanol or saline injections in a separate cohort of male and female mice to identify cell-type specific effects. Cholinergic neurons were identified as positive for *Chat* and negative for *Vglut2* transcripts (*Chat+, Vglut2-*), glutamatergic neurons were positive for *Vglut2* and negative for *Chat* (*Chat-, Vglut2+*), and a third population of co-labeled neurons was identified as expressing both transcripts (*Chat+, Vglut2+*) (**Fig. 3**). We found that the total number of cholinergic neurons was not different between sex and treatment group in the PPN (F_sexXtreatment_(1, 11.5)=0.056, *P*=0.82; F_sex_(1, 11.2)=0.008, *P*=0.93; F_treatment_(1, 11.5)=1.411, *P*=0.26) or in the LDT (F_sexXtreatment_(1, 12.6)=1.521, *P*=0.24; F_sex_(1, 13.0)=1.001, *P*=0.34; F_treatment_(1, 12.6)=0.130, *P*=0.72; **Fig. S2a-b**), which is similar to our results obtained with IHC showing no differences in the number of cholinergic neurons across sex or treatment group. The total number of glutamatergic neurons was also not significantly different between sex and treatment group in the PPN (F_sexXtreatment_(1, 11.6)=1.901, *P*=0.19; F_sex_(1, 11.0)=0.041, *P*=0.84; F_treatment_(1, 11.6)=1.923, *P*=0.19) or in the LDT (F_sexXtreatment_(1, 12.4)=0.223, *P*=0.64; F_sex_(1, 12.4)=0.413, *P*=0.53; F_treatment_(1, 12.4)=1.158, *P*=0.30; **Fig. S2c-d**).

**Figure 3.**
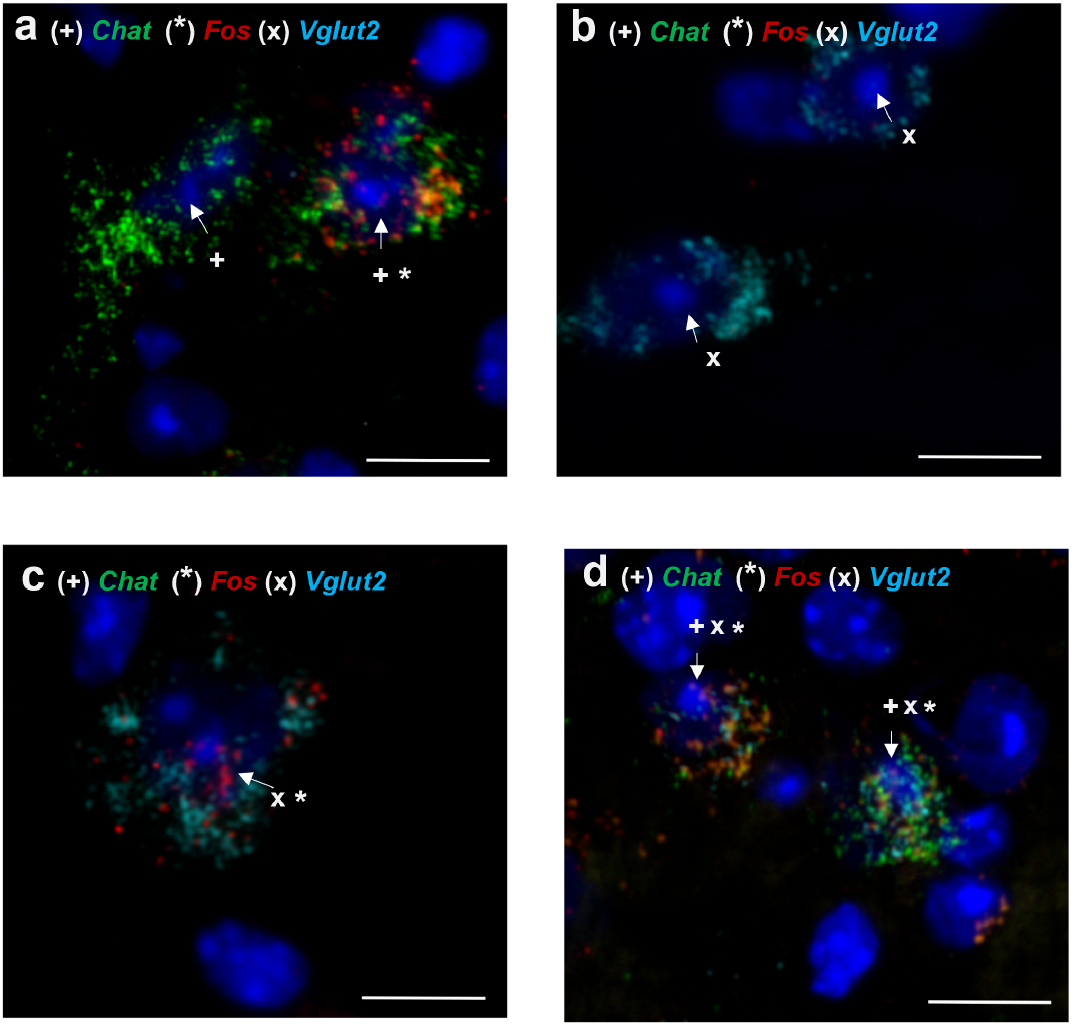
Cholinergic, glutamatergic and co-labelled neurons detected with RNAscope in the MPT. Examples of **a)** *Fos*-positive and negative cholinergic (*Chat*+, *Vglut2-*) neurons, **b)** *Fos*-negative glutamatergic (*Chat*-, *Vglut2+*) neurons, **c)** *Fos*-positive glutamatergic (*Chat*-, *Vglut2+*) neurons and d) *Fos*-positive co-labeled (*Chat*+, *Vglut2+*) neurons. Scale bar = 10 μm.

### Chronic ethanol activation of cholinergic and glutamatergic neuronal cell types in the MPT

We examined the activation of cholinergic, glutamatergic and co-labeled neurons in mice administered 15 daily injections of 2.0 g/kg ethanol or saline. In male mice, we found that the percent of activated cholinergic neurons (*Chat+, Vglut2-, Fos+*) in the PPN was greater in ethanol-treated mice compared with saline-treated mice (nested *t*=3.796, \*\*\**P*=0.0005), with 50.7 ± 2.8% of cholinergic neurons expressing *Fos* in the saline-treated group and 66.8 ± 2.6% in the ethanol-treated group (**Fig. 4a**). We calculated the percent of *Fos* transcript in the activated cholinergic neurons and found a non-significant trend for increased *Fos* transcript levels in the activated cholinergic neurons from ethanol-treated compared with saline-treated mice (nested *t*=1.766, *P*=0.13; saline 2.6 ± 0.3%, ethanol 4.1 ± 0.3%, **Fig. 4b**). This data is consistent with our findings with IHC showing that 15 days of daily 2.0 g/kg ethanol injections increased the percent of activated cholinergic neurons in the PPN. Chronic ethanol treatment also increased the percent of activated glutamatergic neurons (*Chat-, Vglut2+, Fos+*) in the PPN in males (nested *t*=4.154, \*\**P*=0.006, **Fig. 4e**), with 41.7 ± 1.9% of glutamatergic neurons expressing *Fos* in the saline-treated group and 63.1 ± 1.9% of glutamatergic neurons expressing *Fos* in the ethanol-treated group, indicating that chronic ethanol activates both cholinergic and glutamatergic neurons in the PPN in male mice. We found a trend for increased amount of *Fos* transcript in the activated glutamatergic neurons from ethanol-treated mice compared with saline-treated mice (nested *t*=2.039, *P*=0.09, saline 2.2 ± 0.2%, ethanol 3.3 ± 0.2%, **Fig. 4f**). In the LDT in male mice, there was no significant increase in the percent of activated cholinergic neurons (nested *t*=0.858, *P*=0.42; saline 46.4 ± 5.6%, ethanol 56.9 ± 5.5%, **Fig. 4c**) or in the percent of activated glutamatergic neurons (nested *t*=0.846, *P*=0.44; saline 49.7 ± 3.4%, ethanol 54.6 ± 3.1%, **Fig. 4g**) after chronic treatment. Within the activated cholinergic and glutamatergic neurons, there was no difference in the amount of *Fos* transcript expressed across ethanol-or saline-treated groups (cholinergic neurons: nested *t*=0.429, *P*=0.68, saline 3.5 ± 0.5%, ethanol 4.0 ± 0.7%; glutamatergic neurons: nested *t*=1.050, *P*=0.33, saline 2.9 ± 0.4%, ethanol 3.8 ± 0.5%, **Fig. 4d,h**). These data are consistent with our results obtained with IHC which showed no increase in activated cholinergic neurons in the LDT in male mice.

**Figure 4.**
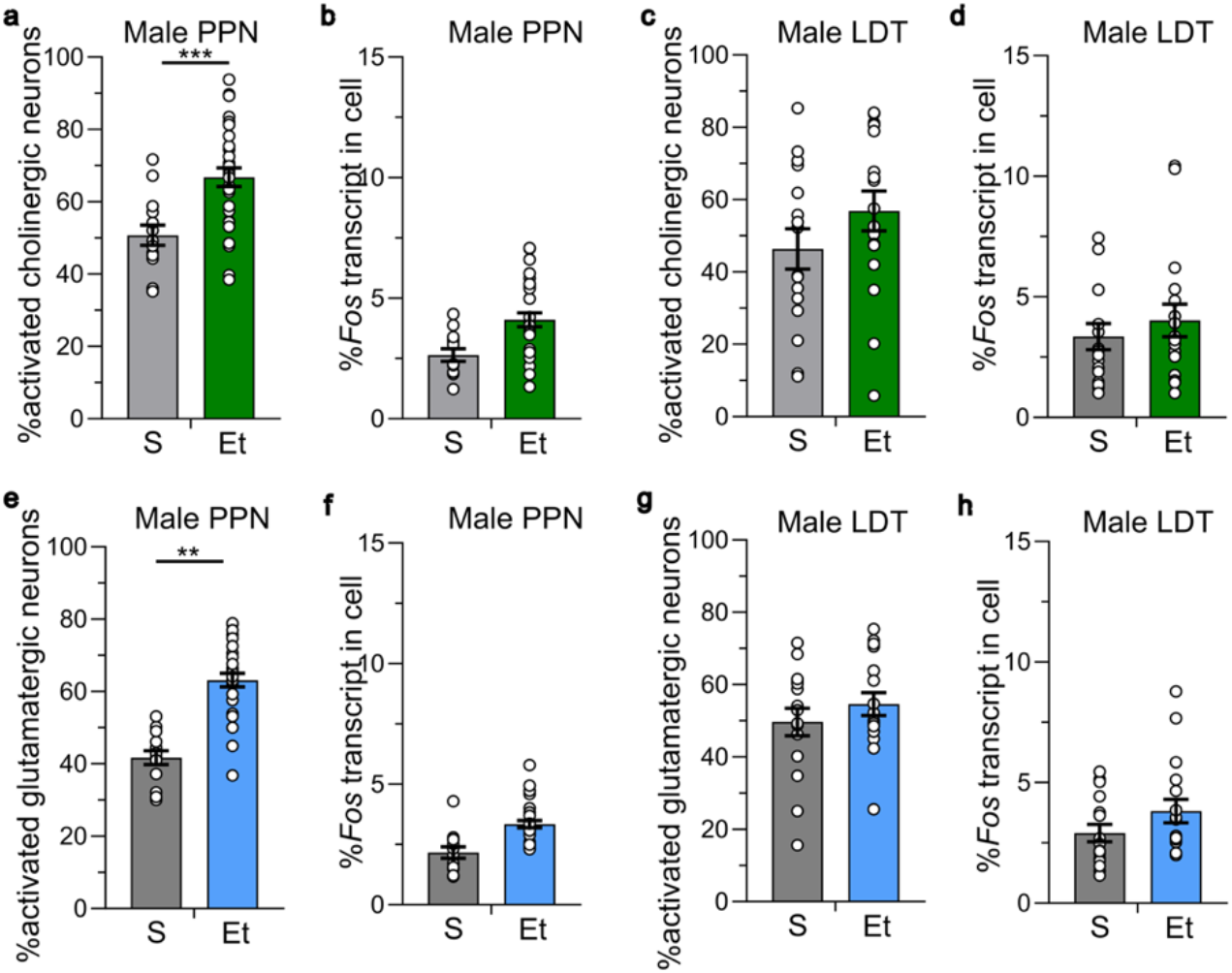
Chronic ethanol activates cholinergic and glutamatergic neurons in the MPT in male mice. Ethanol (2.0 g/kg for 15 days) **a)** increases the percent of activated cholinergic (*Chat*+, *Vglut2-*) neurons in the PPN measured with RNAscope. \*\*\**P*=0.0005, saline (S) vs ethanol (Et) treated groups. **b)** The percent amount of *Fos* transcript in activated neurons in the saline- and ethanol-treated groups was not different. **c-d)** Chronic ethanol did not increase the percent of activated cholinergic LDT neurons or the percent of *Fos* expression in activated neurons in the LDT. **e)** Chronic ethanol increased the percent of activated glutamatergic (*Chat*-, *Vglut2+*) neurons in the PPN. \*\**P*=0.006, S vs Et. **f)** There was no difference in the percent of *Fos* expression in activated neurons between groups. **g-h)** Chronic ethanol did not increase the percent of activated glutamatergic LDT neurons or alter the amount of *Fos* expression in activated neurons in the LDT. PPN: *n*=14-30 ROI from 3-5 mice/group. LDT: *n*=16-17 ROI from 3-5 mice/group. Data expressed as mean ± SEM.

In female mice, the percent of activated cholinergic neurons (*Chat+, Vglut2-, Fos+*) in the PPN and LDT were not significantly different between the saline- and ethanol-treated groups (PPN: nested *t*=0.414, *P*=0.69, saline 55.8 ± 2.9%, ethanol 51.6 ± 5.3%, **Fig. 5a**; LDT: nested *t*=0.069, *P*=0.95, saline 49.2 ± 5.5%, ethanol 50.2 ± 4.8%, **Fig. 5c**). There was also no difference in the amount of *Fos* transcript expressed in activated cholinergic neurons between saline- and ethanol-treated groups in either subregion (PPN: nested *t*=0.497, *P*=0.63, saline 3.6 ± 0.4%, ethanol 3.0 ± 0.5%, **Fig.5b;** LDT: nested *t*=0.421, *P*=0.69, saline 3.9 ± 0.8%, ethanol 3.5 ± 0.6%, **Fig. 5d**). There was no difference in the activated glutamatergic neurons between saline-or chronic ethanol-treated groups in the PPN or LDT in female mice (PPN: nested *t*=0.952, *P*=0.37; saline 56.3 ± 2.5%, ethanol 52.0 ± 2.8%, **Fig. 5e**; LDT: nested *t*=1.414, *P*=0.21; saline 49.3 ± 4.0%, ethanol 58.1 ± 2.6%, **Fig. 5g**). There was also no difference in the amount of *Fos* transcript expressed in activated glutamatergic neurons across treatment group (PPN: nested *t*=0.367, *P*=0.56, saline 5.7 ± 0.7%, ethanol 4.4 ± 0.6%, **Fig. 5f**; LDT: nested *t*=0.183, *P*=0.86, saline 5.2 ± 1.1%, ethanol 4.1 ± 0.7%, **Fig. 5h**).

**Figure 5.**
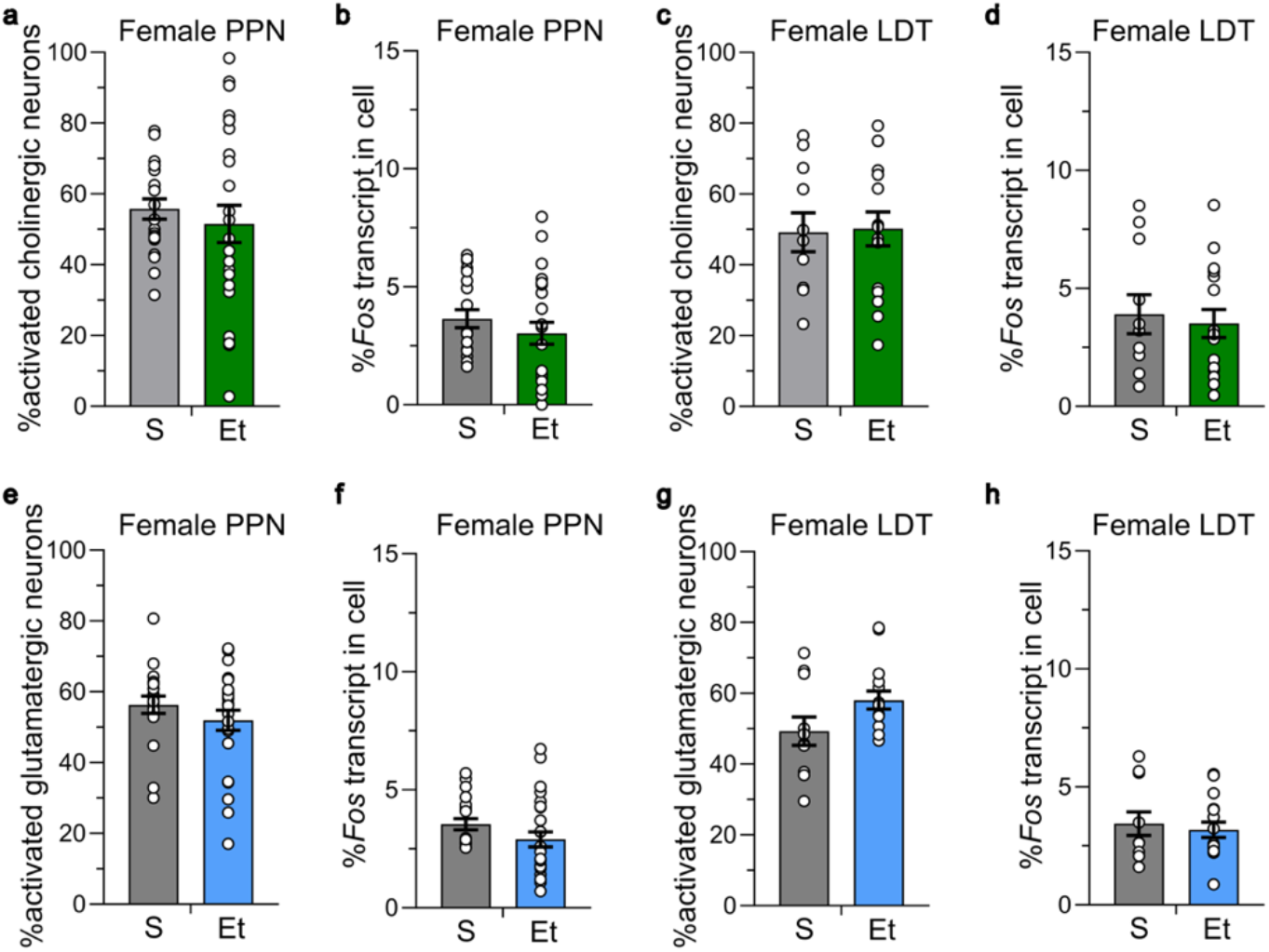
Chronic ethanol did not activate cholinergic and glutamatergic neurons in the MPT in female mice. Ethanol (2.0 g/kg for 15 days) did not increase the percent of activated cholinergic (*Chat*+, *Vglut2-*) neurons in the **a)** PPN or **c)** LDT measured with RNAscope. There was no increase in the percent of activated glutamatergic (*Chat*-, *Vglut2+*) neurons in the **e)** PPN or **g)** LDT. There was no increase in the percent of *Fos* expression in activated **b,d)** cholinergic neurons or in the **f, h)** glutamatergic neurons in the PPN and LDT. PPN: *n*=20-26 ROI from 4-5 mice/group. S=saline, Et=ethanol. LDT: *n*=11-16 ROI from 4-5 mice/group. Data expressed as mean ± SEM.

### Constitutive activation of cholinergic and glutamatergic co-labelled neurons

The total number of co-labeled cholinergic and glutamatergic neurons was not different across sex or treatment group in the PPN (F_sexXtreatment_(1, 13.3)=0.009, *P*=0.93; F_sex_(1, 12.9)=0.064, *P*=0.80; F_treatment_(1, 13.3)=0.722, *P*=0.41) or the LDT (F_sexXtreatment_(1, 12.8)=0.357, *P*=0.56; F_sex_(1, 13.0)=0.0001, *P*=0.99; F_treatment_(1, 12.8)=0.125, *P*=0.73, **Fig. S3**). This co-labeled population was approximately one quarter of all cholinergic neurons, with 27.1 ± 1.7% of the cholinergic neurons in the PPN and 21.5 ± 3.2% of the cholinergic neurons in the LDT also expressing *Vglut2* in male mice. In female mice, 27.1 ± 3.1% of cholinergic neurons in the PPN and 26.1 ± 6.3% of the cholinergic neurons in the LDT also expressed *Vglut2*. The co-labeled population was approximately one tenth of all glutamatergic neurons in the MPT, with 11.5 ± 1.4% of glutamatergic neurons in the PPN and 12.2 ± 2.1% of glutamatergic neurons in the LDT also expressing *Chat* in male mice. In female mice, 10.6 ± 2.3% of glutamatergic neurons in the PPN and 9.8 ± 2.9% of glutamatergic neurons in the LDT also expressed *Chat*. These data indicate the number of co-labeled neurons in the MPT is not different across sex or treatment group.

The cholinergic and glutamatergic co-labeled population in the PPN and LDT in males had a much higher level of activation that occurred in both the saline- and ethanol-treated groups. In the PPN, 87.7 ± 1.4% of co-labeled neurons were activated in the saline-treated group and 88.3 ± 2.4% of co-labeled neurons were activated in the ethanol-treated group (nested *t*=0.184, *P*=0.86, **Fig. 6a**). There was a trend for activated co-labeled neurons from ethanol-treated mice to have a higher percent *Fos* transcript compared with saline-treated mice in the PPN (nested *t*=2.160, *P*=0.07; saline 3.8 ± 0.4%, ethanol 5.7 ± 0.4%, **Fig. 6b**). In the LDT, 87.2 ± 2.5% of co-labeled neurons were activated in the saline-treated group and 93.8 ± 1.6% of co-labeled neurons were activated in the ethanol-treated group (nested *t*=1.898, *P*=0.12, **Fig. 6c**). There was no significant difference in amount of *Fos* transcript expressed in activated neurons across groups (nested *t*=0.748, *P*=0.48; saline 5.0 ± 0.8%, ethanol 6.2 ± 0.9%, **Fig. 6d**).

**Figure 6.**
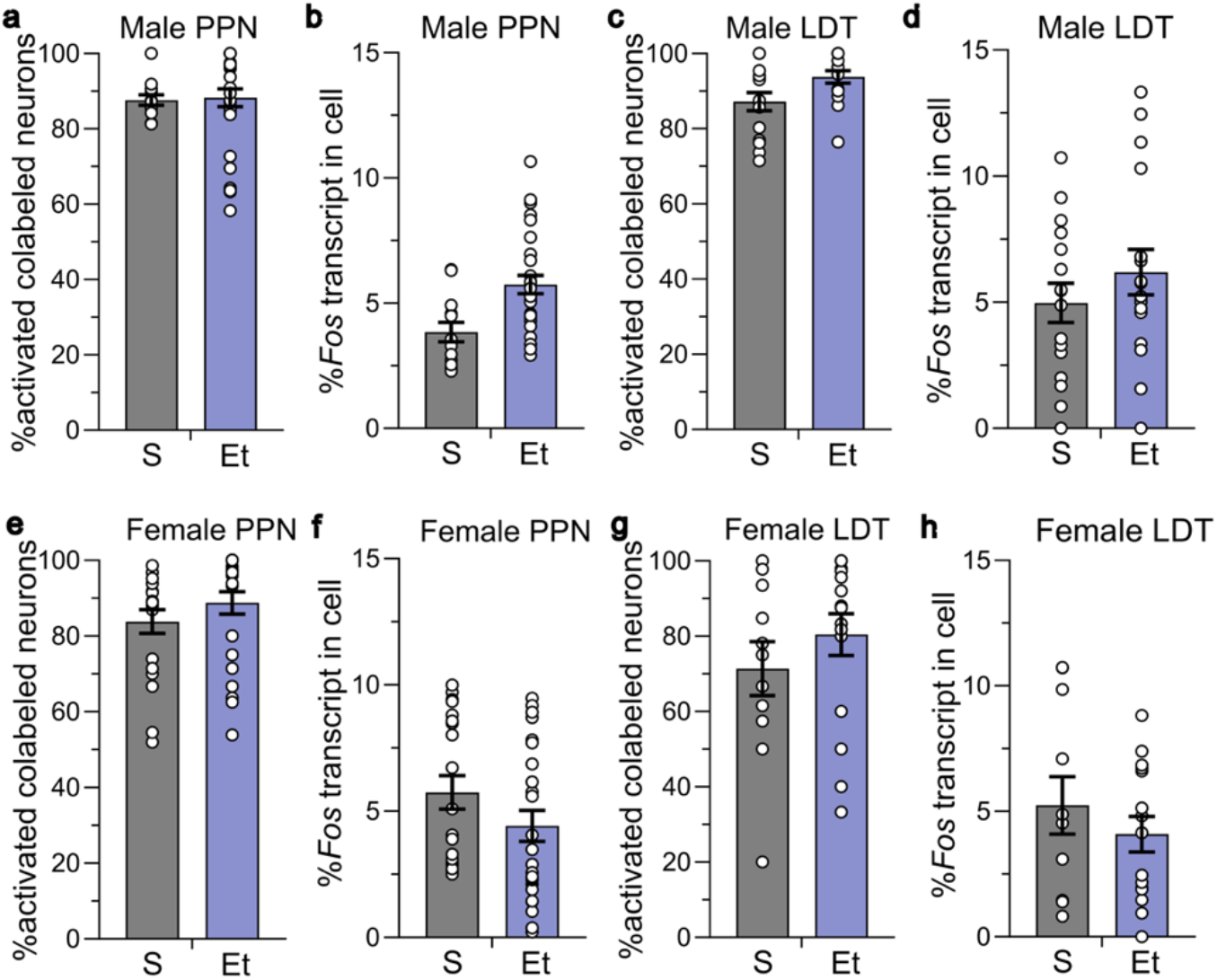
Co-labelled cholinergic and glutamatergic neurons in the MPT had high levels of activation. Ethanol (2.0 g/kg for 15 days) did not increase the percent activation of co-labeled cholinergic and glutamatergic (*Chat*+, *Vglut2+*) neurons. Co-labeled neurons in male **a)** PPN and **c)** LDT, and in female **e)** PPN and **g)** LDT that were active in the S- and Et-treated groups. There was no difference in the percent of *Fos* expression in activated co-labeled neurons in **b,d)** male or **f, h)** female mice. ROI and animal numbers are listed in figure legends 4 and 5. Data expressed as mean ± SEM.

The cholinergic and glutamatergic co-labeled population in the PPN and LDT in female mice also showed a much higher activation level that occurred in both treatment groups, with no significant difference between groups. In the PPN, 83.8 ± 3.1% of co-labeled neurons were activated in the saline-treated group and 88.8 ± 3.0% were activated in the ethanol-treated group (nested *t*=0.897, *P*=0.40, **Fig. 6e**). In the LDT, 71.4 ± 7.2% of co-labeled neurons were activated in the saline-treated group and 80.4 ± 5.6% were activated in the ethanol-treated group (nested *t*=0.502, *P*=0.63, **Fig. 6g**). There was no difference in the percent of *Fos* transcript expressed in activated co-labeled neurons in the saline- and ethanol-treated groups in both subregions (PPN: nested *t*=0.367, *P*=0.56, saline 5.7 ± 0.7%, ethanol 4.4 ± 0.6%, **Fig. 6g**; LDT: nested *t*=0.183, *P*=0.86, saline 5.2 ± 1.1%, ethanol 4.1 ± 0.7%, **Fig. 6h**). These data show that the co-labeled cholinergic and glutamatergic neuronal population in both males and females are highly active regardless of treatment group.

## Discussion

We found that ethanol-induced activation of MPT neurons was dependent on sex, MPT subregion, ethanol dose and time course. Male mice showed activation of cholinergic neurons in the PPN, but not the LDT, after an acute 4.0 g/kg ethanol dose, which is a sedating and aversive dose in mice [31], but not after an acute 2.0 g/kg ethanol dose, which is a dose that can produce conditioned place preference (CPP) in mice [30]. Cholinergic and glutamatergic neurons of the PPN, but not the LDT, were also activated after 15 days of daily 2.0 g/kg ethanol injections in male mice. In contrast, female mice showed no activation of cholinergic or glutamatergic neurons in the PPN or LDT after the same acute or chronic ethanol injections Lastly, we found a population of co-labeled neurons expressing both cholinergic and glutamatergic cell markers that were highly active in both the saline and ethanol treatment conditions in both sexes. Overall, our data show that the brainstem MPT response to ethanol is dramatically different by sex and by MPT subregion, which suggests that cholinergic and glutamatergic signaling is differentially activated by ethanol. These data highlight the importance of investigating both sexes and has important implications on design of pharmacological therapies that are based on male neurobiology only.

The MPT has previously been implicated in the effects of misused drugs. Prior studies have shown that a single systemic injection of nicotine at a dose of 0.3 or 1.0 mg/kg increased c-Fos expression in the PPN and LDT of male rats measured with IHC [19], although which neuronal cell types were activated was not determined. Male rats that received twice daily injections of 10 mg/kg morphine for 6.5 days also showed more intense staining for vesicular ACh transporter protein expression in the LDT [32]. In this study, we investigated the effect of acute and chronic ethanol on the activation of MPT neurons in male and female mice. We found that acute administration of 4.0 g/kg ethanol and chronic administration of 2.0 g/kg ethanol, increases c-Fos protein in cholinergic PPN neurons, and increases *Fos* transcript in cholinergic and glutamatergic PPN neurons, respectively. Past studies have shown that ethanol consumption increases ACh levels in the VTA, and the only source of ACh to the VTA originates in the MPT [7,11,12]. The ethanol-induced activation of the PPN suggests that this subregion may be important in modulating ethanol-related behaviors. Inactivating PPN neurons in a non-cell-type specific manner with infusions of GABA agonists baclofen and muscimol has been shown to reduce voluntary nicotine intake in male rats [21,33]; however, the PPN may not mediate the actions of all misused drugs as lesioning PPN cholinergic neurons did not affect cocaine or heroin self-administration or attenuate cocaine or heroin CPP in male rats [34]. Intriguingly, in male mice made physically dependent on ethanol using the Lieber DeCarli diet, lesioning the PPN attenuated the expression of ethanol CPP after conditioning with a low dose of ethanol (0.2 g/kg *i.p.*); however, lesioning the PPN did not attenuate ethanol CPP in non-dependent mice [35], suggesting the PPN may be differentially modulating ethanol responses based on ethanol exposure levels. In our study, we did not examine physical withdrawal signs in mice chronically treated with 2.0 g/kg ethanol, thus it is unclear if the activation of cholinergic and glutamatergic neurons after 15 days, but not after 1 or 5 days, of ethanol injections coincided with the development of physical dependence. Our data further implicate cholinergic and glutamatergic PPN neurons in drug-related effects and we speculate that the PPN is recruited after chronic administration of ethanol and may mediate ethanol-related behaviors.

We found no increase in c-Fos protein or *Fos* transcript in the LDT in either cholinergic or glutamatergic neurons in male mice, highlighting a MPT subregion difference in ethanol responsiveness. PPN and LDT cholinergic neurons have overlapping inputs and projection targets; however, recent studies focusing on MPT cholinergic inputs to the VTA and substantia nigra have begun to distinguish complex and subtle functional differences between the two MPT subregions. In male rats, a greater proportion of cholinergic neurons from the PPN project to the substantia nigra, whereas a greater proportion of LDT cholinergic neurons project to the VTA [36]. The LDT also contains more cholinergic neurons that project to non-VTA and non-substantia nigra locations compared with the PPN [36]. Stimulation of LDT cholinergic axons in the VTA preferentially caused activation of VTA dopamine neurons that project to the nucleus accumbens, whereas stimulation of PPN cholinergic axons in the VTA primarily activated dopamine neurons that do not project to the accumbens [11]. However, other studies have shown that optogenetic stimulation of both PPN and LDT cholinergic terminals in the VTA promotes CPP [36], indicating some functional overlap between the two subregions. Lastly, LDT cholinergic neurons project to the striatum and modulate cholinergic interneurons, as inhibiting LDT, but not PPN, cholinergic neuron terminal activity in the striatum interferes with goal-directed outcome devaluation [37]. Based on this prior data, we expected ethanol to activate cholinergic neurons in the LDT as this subregion innervates the VTA to a greater extent than PPN, and innervates dopamine neurons that project to the striatum, which are critically important for drug-related behaviors [11,36]. However, we found that acute and chronic ethanol activated PPN, but not LDT, cholinergic neurons in male mice, suggesting that ethanol selectively recruited PPN cholinergic neuron activity. Determining the mechanisms underlying the specific recruitment of PPN, and not LDT, neurons by ethanol will be critical to better understand the neurobiology of ethanol. It is likely that the differences in function between the PPN and LDT in projection targets outside of the VTA and substantia nigra, which have not been well studied, will also be important in understanding this subregion specificity.

We found that chronic ethanol administration induced activation of PPN glutamatergic neurons in male mice. Unlike the bouton neurotransmission of cholinergic neurons, glutamatergic neurons have direct innervation targets, providing excitatory drive to post-synaptic neurons. Glutamatergic neurons have been shown to be intermingled with the cholinergic population in the MPT in male rats [14], and we also observe intermingling of these two neuronal populations in both male and female mice. MPT glutamatergic neurons have more recently been implicated in drug-related actions, as optogenetic inhibition of LDT glutamatergic neurons or glutamatergic afferents from the LDT to the VTA impairs the development of cocaine locomotor sensitization in male mice [38]. In addition, optogenetic activation of PPN glutamatergic neurons can activate VTA dopamine and non-dopamine neurons, and can support self-stimulation behaviors [15]. We found that chronic ethanol induced the activation of glutamatergic neurons in the PPN, and not the LDT, which mirrors the PPN-specific activation of cholinergic neurons in male mice. This indicates that ethanol-induced selective activation of the PPN involves multiple neuronal cell types. Identification of the projection targets of these activated glutamatergic neurons would provide additional insight into the neural circuits that are activated by ethanol.

Notably, almost all the past studies on misused drugs and the MPT were conducted in male animals and there is little to no data in female animals. We found a striking sex difference in the responsiveness of cholinergic and glutamatergic neurons to ethanol in the MPT as we did not see increases in the activation of cholinergic or glutamatergic neurons in either the PPN or LDT in female mice. Moreover, in the PPN the baseline level of neuronal activation in cholinergic neurons was significantly different in male and female mice, with female mice having a higher level of cholinergic cell activation. We have previously observed sex differences in cholinergic mechanisms in mice, as we found that the *Chrna6* and *Chrnb3* nAChR gene cluster was differentially regulated by sex [29,39]. There are also important sex differences in ethanol consumption and ethanol-related behaviors in mice, as female C57BL/6J mice consume more ethanol compared with male C57BL/6J mice [40–42]. We have also observed sex differences in compensatory behavior after quinine-adulteration of ethanol, as a three-bottle choice test showed that female mice presented with increasing concentrations of quinine in the ethanol bottle respond by consuming more nicotine solution, whereas male mice consume more water [43]. Prior work shows that ethanol clearance after a single *i.p.* injection of 3 g/kg is not significantly different between male and female C57BL/6J mice [44], suggesting that these changes in ethanol effects and consumption are likely not due to sex differences in ethanol clearance.

We did not track estrus cycle in our female mice in this study or in our prior work examining chronic ethanol consumption [40,43], and estrus cycle does not affect voluntary ethanol consumption levels in mice [45]. However, sex hormones may still have an important role in regulating ethanol effects as estrogen receptors have been shown to interact with glutamatergic signaling and affect VTA neuronal activity [46]. Investigating sex differences in cholinergic mechanisms in rodents and humans is necessary to understand the sex differences in AUD, as women have a faster progression to ethanol dependence and more detrimental outcomes compared with men [47–50]. Moreover, varenicline, a nAChR partial agonist, is in clinical trials for AUD and is more effective in men compared with women [51], highlighting the need to understand sex differences in cholinergic mechanisms to improve rationale drug design efforts for AUD that will be effective in both men and women.

We found a population of neurons co-labeled with cholinergic and glutamatergic markers in both male and female mice, suggesting that these neurons release both ACh and glutamate. Prior work examining the MPT in male rats showed approximately 5% of co-labeled cholinergic and glutamatergic neurons [14]. Here, we found that co-labeled neurons were approximately one-quarter of the total cholinergic neuron population in both the PPN and LDT, in both males and female mice. It is possible that a species difference underlies the difference in the proportion of co-labeled neurons. Intriguingly, most co-labeled neurons were active in the saline- and ethanol-treated conditions, suggesting that this population is functionally distinct from cholinergic and glutamatergic only neurons. The MPT is also implicated in fundamental behaviors such as sleep and movement, and is a target for deep brain stimulation for the treatment of Parkinson’s disease [52]. The co-labeled population may have a larger contribution to these roles, which would be active in the saline- and ethanol-treated groups. Notably, changes in the MPT are implicated in the poor sleep quality observed in AUD [53].

We find that ethanol activates cholinergic and glutamatergic neurons of the PPN and not the LDT in male mice. In contrast, ethanol had no effect on the activation of cholinergic or glutamatergic neurons in the MPT of female mice. Female mice had higher baseline activation of cholinergic neurons compared with males, suggesting that endogenous cholinergic signaling differs by sex. These findings illustrate the differential effects of ethanol across dose, duration, MPT subregion and sex, indicating that the interactions between ethanol and the MPT are more complex than previously thought.

## Acknowledgments

This work was supported by R01 AA026598 (AML), T32 DA007097 (SMM).

## Author Contributions

SMM, EA, SE, KP, CW, LH collected the data, RG wrote the MATLAB code, RG, SE, SMM, AM analyzed the data, SMM and AML wrote and edited the manuscript.

## Data Availability Statement

The datasets generated during and analyzed, and MATLAB code are available from the corresponding author on request.

## Supplementary Figures and Legends

**Figure S1.**
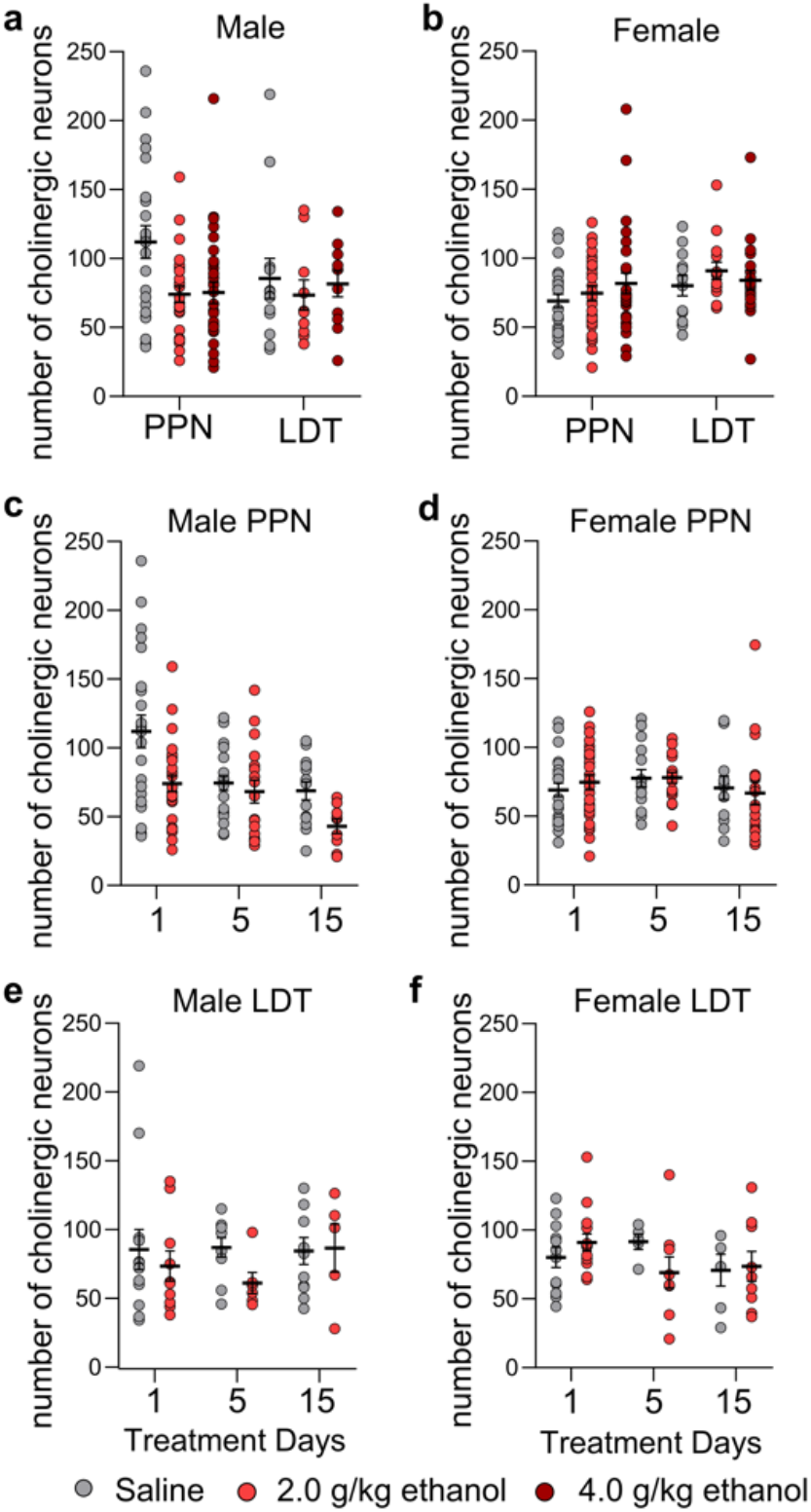
The number of cholinergic neurons identified by IHC in the MPT is not different across sex, MPT subregion or treatment group. The number of cholinergic neurons in **a)** male PPN and LDT, and **b)** female PPN and LDT acutely treated with saline-, 2.0 or 4.0 g/kg ethanol. The number of cholinergic neurons in **c)** male PPN, **d)** female PPN, **e)** male LDT and **f)** female LDT chronically treated with 1, 5 or 15 daily injections of saline or 2.0 g/kg ethanol. The number of ROI and animals per group are listed in figure legend 2. Data expressed as mean ± SEM.

**Figure S2.**
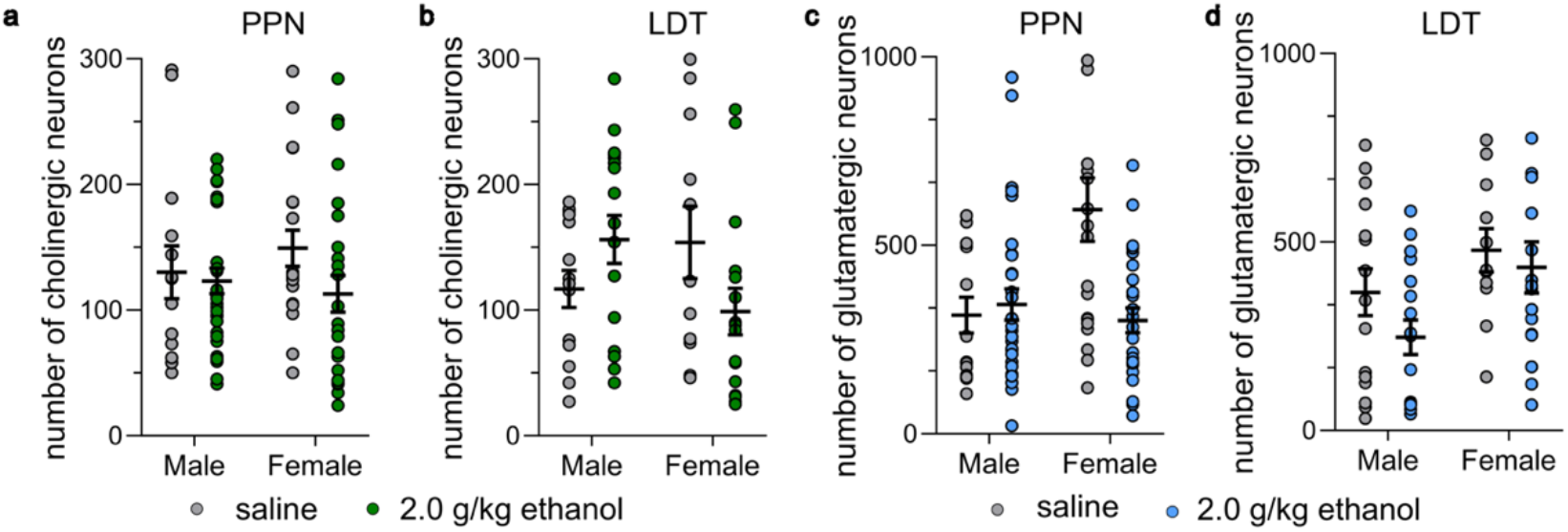
The number of cholinergic and glutamatergic neurons identified by RNAscope in the MPT is not different across sex, MPT subregion or treatment group. The number of cholinergic neurons in the **a)** PPN and **b)** LDT and the number of glutamatergic neurons in the **c)** PPN and **d)** LDT in male and female mice chronically treated with 15 daily injections of saline-or 2.0 g/kg ethanol. The number of ROI and animals per group are listed in figure legends 4 and 5. Data expressed as mean ± SEM.

**Figure S3.**
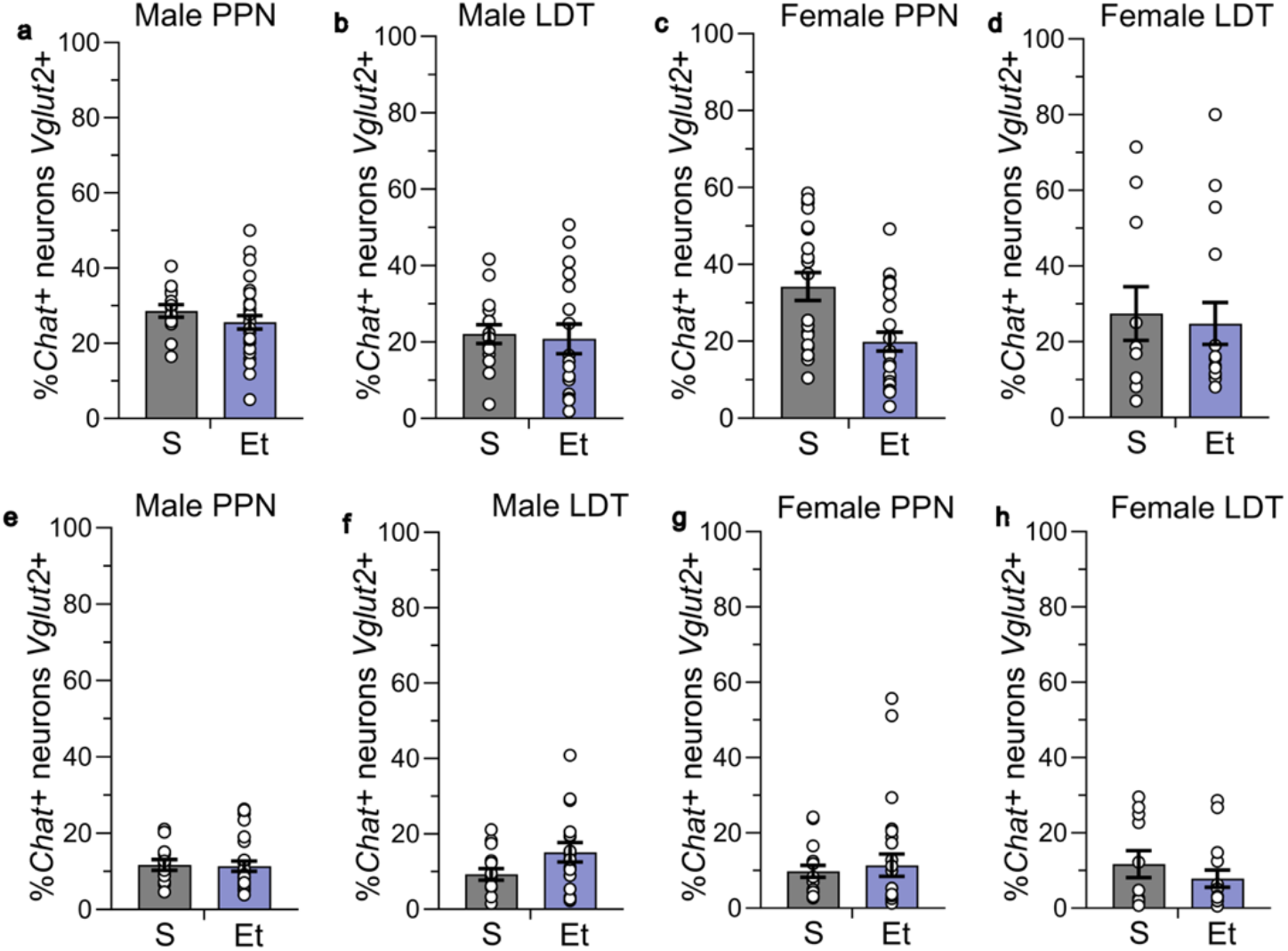
The number of co-labeled cholinergic and glutamatergic neurons identified by RNAscope in the MPT is not different across sex, MPT subregion or treatment group. The percent of *Chat*-positive neurons in the *Vglut2*-positive population is similar between the saline and chronic-ethanol treated groups in the male **a)** PPN and **b)** LDT and female **c)** PPN and **d)** LDT. The percent of *Vglut2*-positive neurons in the *Chat*-positive population is similar between groups in the male **e)** PPN and **f)** LDT and female **g)** PPN and **h)** LDT. The number of ROI and animals per group are listed in figure legends 4 and 5. Data expressed as mean ± SEM.

